# A multivalent fuzzy interface drives reversible COPII coat assembly

**DOI:** 10.1101/2020.04.15.043356

**Authors:** Viktoriya G. Stancheva, Joshua Hutchings, Xiao-Han Li, Balaji Santhanam, M. Madan Babu, Giulia Zanetti, Elizabeth A. Miller

**Affiliations:** MRC Laboratory of Molecular Biology; Institute of Structural and Molecular Biology, Birkbeck College

## Abstract

Protein secretion is initiated at the endoplasmic reticulum by the COPII coat, which self-assembles to form vesicles. Here, we examine the mechanisms by which the outer scaffolding layer of the coat drives local assembly of a structure rigid enough to enforce membrane curvature, yet able to readily disassemble at the Golgi. An intrinsically disordered region in the outer coat protein, Sec31, drives binding with an inner coat layer via multiple distinct interfaces. Interactions are individually dispensable but combinatorially reinforce each other, suggesting coat oligomerization is driven by the cumulative effects of multivalent interactions. Such a multimodal assembly platform could be readily reversed at the Golgi via perturbation of each individual interface. These design principles provide an explanation for how cells build a powerful yet transient scaffold to direct vesicle traffic.

## Introduction

Eukaryotic cells secrete soluble and membrane-bound proteins to serve a variety of essential functions, including nutrient and ion uptake, cell-cell communication, and environmental signalling. Protein transport within the secretory pathway is facilitated by vesicles that are generated by cytoplasmic coat proteins, which simultaneously recruit appropriate cargo and sculpt the donor membrane into spherical structures. Membrane bending requires significant force to overcome the intrinsic rigidity of the lipid bilayer, and coat proteins solve this problem by self-organizing into oligomeric scaffolds that can impose structure on the underlying membrane. The coordination of coat polymerization with cargo recruitment is essential to avoid dead-end formation of empty vesicles, but inclusion of cargo can further increase the membrane bending energy required by the coat (*1*, *2*). The paradox of such coat scaffolding oligomers is that the resultant assembly must be rigid enough to exert significant force, but also fully reversible to allow uncoating and vesicle fusion. In clathrin-mediated endocytosis, force generation is driven in part by the highly interdigitated clathrin triskelion, which requires ATP hydrolysis to disassemble. In contrast, the COPII and COPI coats, which mediate traffic in the early secretory pathway, form simpler scaffolds that can spontaneously disassemble. Here, we sought to address two aspects of COPII coat function: how coat assembly builds dynamic yet stable structures, and how such structures are reversed to prime fusion with the Golgi.

The COPII coat comprises five proteins that self-assemble on the cytosolic face of the endoplasmic reticulum (ER) membrane to drive traffic of nascent secretory proteins (Fig. 1A). COPII assembly is initiated upon GTP binding by the small GTPase, Sar1, which exposes an amphipathic α-helix that embeds shallowly in the membrane (*3*). GTP-bound Sar1 recruits Sec23/Sec24 (*4*), which is the cargo-binding subunit of the coat (*5*, *6*). The Sar1/Sec23/Sec24 “inner coat” complex in turn recruits the “outer coat”, Sec13/Sec31, to drive vesicle formation (*4*, *7*). Sec13/Sec31 tetramers form rods (*8*) that can self-assemble *in vitro* into a cage-like structure (Fig. S1A) that closely matches the size and geometry of vesicles observed by EM (*9*). Sar1/Sec23/Sec24 can also form higher order assemblies (*10*), but Sec13/Sec31 is required for organization of these arrays (*3*). Binding of the inner coat stimulates GTP hydrolysis on Sar1, and the outer coat further accelerates this reaction. The GTP hydrolysis cycle is key to the dynamic assembly and disassembly of the coat, thereby contributing to a dynamically metastable structure. Although the structures of individual coat layers are well-described, how they assemble locally to create rigid structures remains unclear. Moreover, accessory factors that are not integral to the coat, such as Sec16, also facilitate coat assembly via poorly defined mechanisms. Finally, how these stable oligomers are released from the membrane, both to recycle the coat proteins and to expose vesicle fusion machinery, is unknown (Fig. 1A).

**Figure 1.**
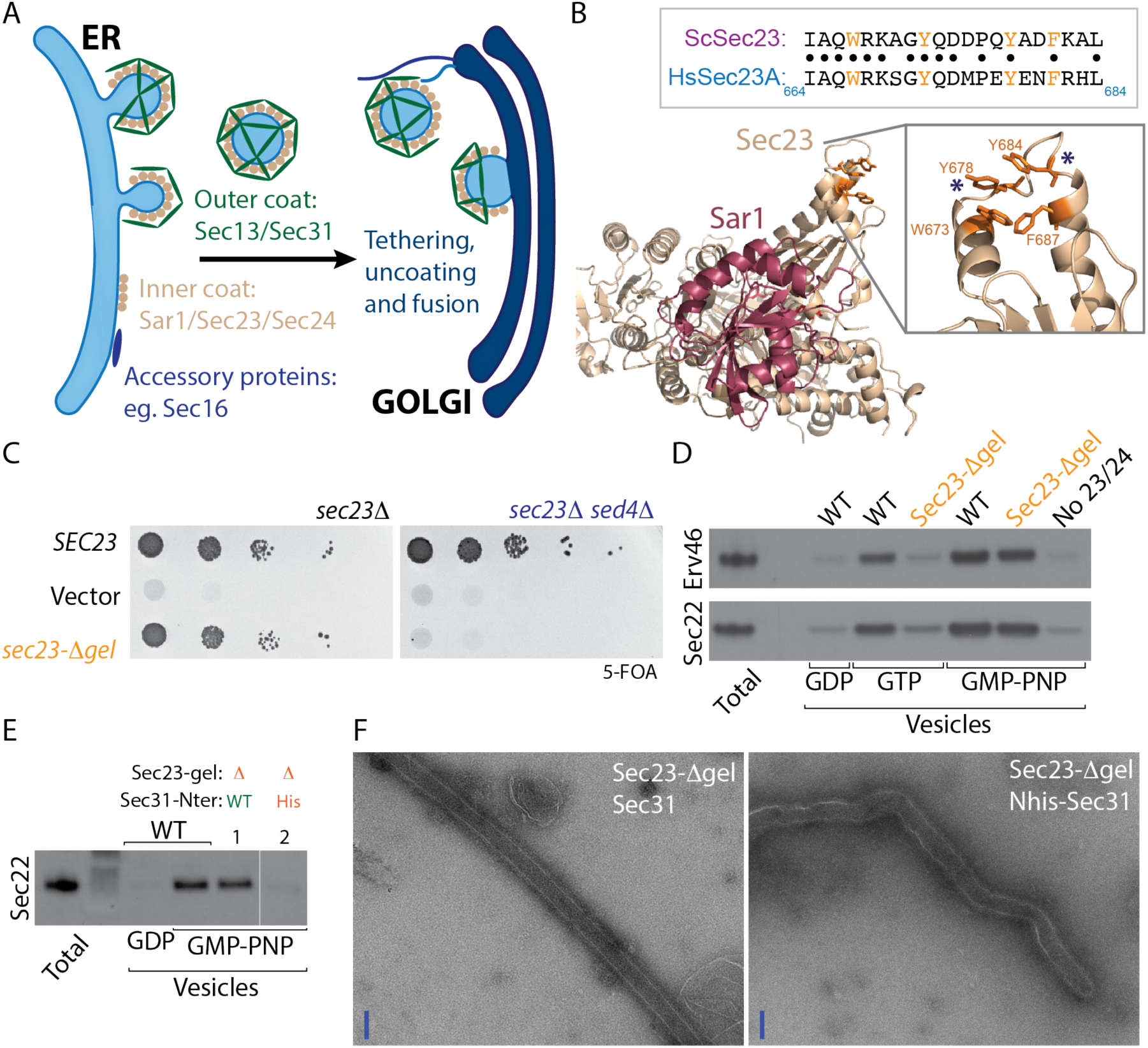
Sec23 gelsolin domain is dispensable for coat assembly but contributes to coat stability. **A.** Cartoon depicting vesicle formation from the endoplasmic reticulum (ER) by the COPII coat, followed by vesicle release, tethering, uncoating and fusion with the Golgi. **B.** Structural model (PDB 1M2O) of Sar1 (red) and Sec23 (wheat) highlighting the membrane-distal gelsolin domain (inset right) that contains conserved aromatic residues (inset left) that in human Sec23 contribute to PPP binding. **C**. *sec23Δ* and *sec23Δ sed4Δ* strains were transformed with the indicated plasmids and growth tested on 5-FOA, which counterselects for a *SEC23::URA* plasmid. Growth on 5-FOA is indicative of ability to serve as the sole copy of *sec23*. Deletion of the gelsolin loop was tolerated in the *sec23Δ* strain but was unable to support viability in the *sec23Δ sed4Δ* double mutant. **D**. *In vitro* budding experiments using yeast microsomal membranes incubated with Sar1, Sec23/Sec24, Sec13/Sec31 and the indicated nucleotides. Vesicle release from the donor membrane is measured by detecting incorporation of COPII cargo proteins, Sec22 and Erv46, into a slowly sedimenting vesicle fraction. When wild-type Sec23 was replaced with Sec23-Δgel, budding was compromised only in the GTP condition. **E**. *In vitro* budding as described for (C) but where one reaction used Sec31 that was tagged with hexahistidine at the N-terminus (lane 2). Sec23-Δgel in combination with Nhis-Sec31 was compromised for budding even in the presence of GMP-PNP. **F.** Negative stain electron microscopy of GUVs incubated with Sar1•GMP-PNP and the indicated wild-type or mutant proteins.

In this study, we aimed to dissect the COPII coat assembly pathway by testing the functional importance of interfaces that drive outer coat recruitment to the inner coat layer and thus trigger membrane scaffolding. Two interactions between the inner and outer coat layers have been characterized biochemically and structurally (*11*, *12*), both of which involve a long, unstructured domain of Sec31 (Fig. S1B). A short “active fragment” of Sec31 lies across the membrane-distal surface of Sec23/Sar1, contacting both proteins and stimulating Sar1 GTPase activity via two key residues, W_922_ and N_923_ in *S. cerevisiae* Sec31 (Fig. S1C) (*11*). A second mode of interaction involves triple-proline (PPP) motifs on Sec31 that bind to the gelsolin domain of Sec23 (*3*, *12*)(Fig. 1B, S1B). Exactly how these inter-subunit interfaces contribute to coat polymerization remains unclear. Moreover, these interfaces likely constitute important platforms for regulatory accessory factors that might modulate the COPII coat during different stages of vesicular transport (*12*, *13*). A number of PPP-containing proteins interact with COPII, for example to aid assembly (Sec16), or to promote formation of large carriers for transport of bulky cargoes in mammals (TANGO1). Here, we combine genetic perturbation with *in vitro* reconstitution assays to test the essentiality of individual interactions and determine the precise stage at which specific mutants are defective. We find that coat assembly is driven by a combination of evolutionarily conserved multivalent interactions between the inner and outer coat layers that combinatorially reinforce each other. Moreover, outer coat oligomerization via a known Sec31-Sec31 structural interface reinforces these interactions, suggesting a feed-forward mechanism for coat propagation. Our data define an assembly pathway that can be modulated by regulatory factors to confer reversibility to a metastable oligomeric structure without the input of energy to drive disassembly. Our findings also suggest a partial mechanism by which cells ensure directionality of vesicle traffic by local coat disassembly triggered at the Golgi by proteins competing for the same interaction interface.

### PPP-driven interactions are dispensable for coat assembly but contribute to coat stability

We first tested the importance of PPP motifs in COPII assembly by mutagenesis of the Sec23 gelsolin domain that is known to interact with PPP motifs. Four critical aromatic residues that form the PPP-binding cleft are conserved between yeast and humans (Fig. 1B), and mutation of these residues abrogated PPP binding to human Sec23A (*12*). We engineered a mutant that replaced key hydrophobic residues in the PPP-binding cleft with a glycine-serine-glycine tripeptide. This gelsolin loop mutant (*sec23-Δgel*) complemented a *sec23Δ* null strain, revealing that PPP-binding is not essential for coat function (Fig. 1C). We next tested whether perturbations to the GTPase cycle of the coat might sensitize yeast to the loss of this PPP-binding interface. Sed4 is a non-essential accessory factor that is thought to assist Sec16 in Sar1 GTP regulation (*13*, *14*). Indeed, *sec23-Δgel* was inviable when *SED4* was also deleted, suggesting that compromised PPP-binding by Sec23 becomes problematic when the GTP cycle of the coat is altered (Fig. 1C). To gain further insight into the nature of the defect associated with perturbation of the gelsolin domain, we used an *in vitro* assay that reconstitutes vesicle formation from purified microsomal membranes (*15*). Vesicle release is monitored by the presence of cargo proteins, Erv46 and Sec22, in a slowly sedimenting vesicle fraction following incubation with purified COPII proteins. This budding assay showed that Sec23-Δgel could drive vesicle formation with GMP-PNP but not with GTP (Fig. 1D). Thus, when coat assembly is stabilized by a non-hydrolysable GTP analog, perturbation of the gelsolin-PPP interaction has minimal effect, but under the condition of GTP-dependent coat turnover (*7*), loss of this interface impairs vesicle formation.

A GTP-specific vesicle budding defect was reminiscent of that observed with N-terminally tagged Sec31, where the histidine tag is thought to interfere with assembly of the cage “vertex” interactions (Fig. S1A) that drive outer coat oligomerization (*3*). In the context of coat turnover promoted by GTP hydrolysis, further destabilization of the coat at cage vertices is incompatible with full coat assembly (*3*). We therefore tested the Sec23-Δgel mutant in the budding assay in the presence of Nhis-Sec31, finding that vesicles fail to form even with GMP-PNP to stabilize the coat (Fig. 1E). Consistent with this, in a membrane bending assay, Sec23-Δgel promoted tubulation of giant unilamellar vesicles (GUVs) into straight lattice-coated tubules in the presence of GMP-PNP and wild type Sec31. However, when GUV tubulation was induced using Sec23-Δgel and Nhis-Sec31, tubes were irregular and no extended coat lattice could be visualised (Fig. 1F). One interpretation of these findings is that two separate interactions – Sec23/Sec31 binding via PPP motifs and cage assembly via Sec31/Sec31 vertex interfaces – mutually reinforce each other to stabilize the coat and propagate coat assembly. Such a feed-forward loop to drive vesicle formation might prevent inappropriate assembly away from the ER membrane. These mutually stabilizing interactions are especially important under conditions of GTP hydrolysis, where the coat is inherently unstable (*7*). Our Sec23 mutagenesis experiments suggest that PPP binding contributes to but is not essential for coat assembly. The importance of the PPP interaction is revealed when the GTP cycle of the coat is perturbed, suggesting that the nucleotide-associated interface and PPP binding are mutually productive in COPII assembly.

We next sought to test the importance of specific PPP motifs by mutagenesis of Sec31, additionally abrogating the “active fragment” that also contributes to inner-outer coat binding (Fig. 2A). Within the active fragment, residues W_922_ and N_923_ drive GTPase stimulation (*11*) and are likely the most stable part of the interaction (*3*). Alanine substitution of these amino acids had no effect on cell viability (Fig. 2B). Similarly, glycine/serine substitution of all 6 PPP motifs, plus an additional PPAP motif that might also bind to the gelsolin domain (*12*) (*sec31-ΔPPP*) caused no growth defect, even when combined with the catalytic W_922_A/N_923_A mutation (Fig. 2B). In a coat recruitment assay that measures binding of purified proteins to large (400nm) unilamellar liposomes in the presence of GMP-PNP, all variant proteins were recruited normally (Fig. S2A). Moreover, when the different mutants were used in microsomal budding reactions, each protein supported vesicle formation both with GTP and GMP-PNP, although vesicle formation was reduced with the combined W_922_A/N_923_ A/ΔPPP mutant (Fig. 2C). We tested whether destabilization of cage interfaces by the introduction of the N-terminal His tag would sensitize these mutants as it had for the *sec23-Δgel* mutant. Although Nhis-Sec31 supports viability, in this background the ΔPPP and W_922_A/N_923_A/ΔPPP mutants were inviable (Fig. 2D). Nonetheless, these mutant proteins were competent in the coat recruitment (Fig. 2E, Fig. S2B) and tubulation assays (Fig. 2F). Thus, the known interaction motifs cannot explain the entirety of inner coat/outer coat assembly, nor of inner coat array formation.

**Figure 2.**
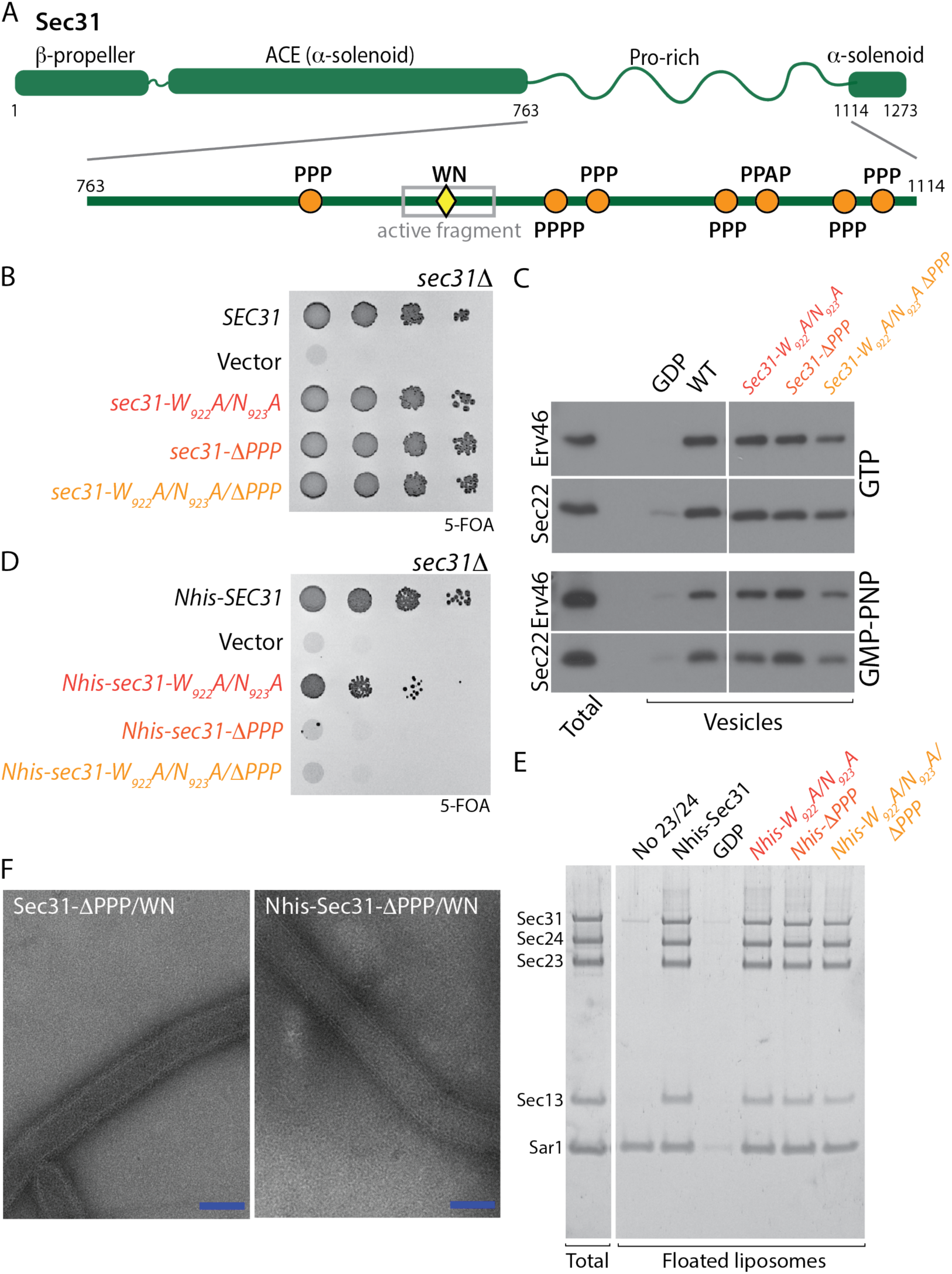
Sec31 known interaction interfaces are individually dispensable for coat assembly. **A.** Diagram of Sec31 showing structured elements (β-propeller and α-solenoids) and the proline-rich disordered region that contains the active fragment (grey box) with catalytic W_922_/N_923_ residues (yellow diamond) and PPP motifs (orange circles) indicated. **B**. Serial dilutions of *sec31Δ* cells transformed with the plasmids indicated spotted onto 5FOA; mutation of the catalytic residues and/or the PPP motifs had no effect on viability. **C.** *In vitro* budding from yeast microsomes with the proteins indicated revealed normal budding efficiency, except for the Sec31-W_922_A/N_923_/ΔPPP mutant, which was reduced in its ability to generate vesicles. **D.** Serial dilutions of a *sec31Δ* strain expressing Nhis-Sec31 mutants as indicated were grown on 5FOA, revealing lethality associated with the combination of the N-terminal His tag and loss of PPP motifs. **E.** Purified COPII proteins (indicated on left in the total fraction) were incubated with synthetic small unilamellar liposomes and floated on a sucrose gradient. Buoyant liposomes and bound proteins were collected and analysed by SDS-PAGE and SYPRO Ruby staining. All mutant protein variants were recruited to liposomes with equal efficiency as the wild-type protein. **F**. Negative stain electron microscopy of GUVs incubated with the indicated proteins.

### A charge-driven interaction interface contributes to coat assembly

Since the catalytic and PPP motifs didn’t fully account for inner/outer coat interactions, we examined the Sec31 disordered region for additional features that might contribute to coat assembly (*16*). We calculated charge properties across Sec31, focusing on the fraction of charged residues and net charge per residue. We found extensively charged clusters of residues across different regions of Sec31, including clusters of net negatively charged residues in the structured domain, and smaller clusters of net positive charges in the disordered region (Fig. 3A). These positive charge clusters were also features of Sec31 orthologues in human and Arabidopsis (Fig. S3A), suggestive of an evolutionarily conserved property. We searched for a corresponding surface on Sec23 that may comprise a charge-driven interaction interface between the inner and outer coats. We identified a negatively charged surface adjacent to the gelsolin domain, separate from that occupied by the “active fragment” (Fig. 3B, top panel). Reversing the charges of this region on Sec23 (Fig. 3B, bottom panel) led to lethality (Fig. 3C), and the purified protein was defective for recruitment of Sec13/Sec31 in the liposome binding assay (Fig. 3D). We propose that charge-based interactions between Sec23 and Sec31 represent a third binding mode that contributes to inner/outer coat interactions (Fig. 3E(i)). We note that three different types of interaction drive association of the inner and outer coats: a nucleotide-driven interface mediated by W/N residues, linear (PPP) motifs, and a charge-driven interaction. Whether the W/N, PPP and charge interactions drive binding to the same Sec23 molecule, or instead bridge adjacent Sec23/Sec24 complexes to contribute to inner coat assembly remains to be determined (Fig. 3E(i)).

**Figure 3.**
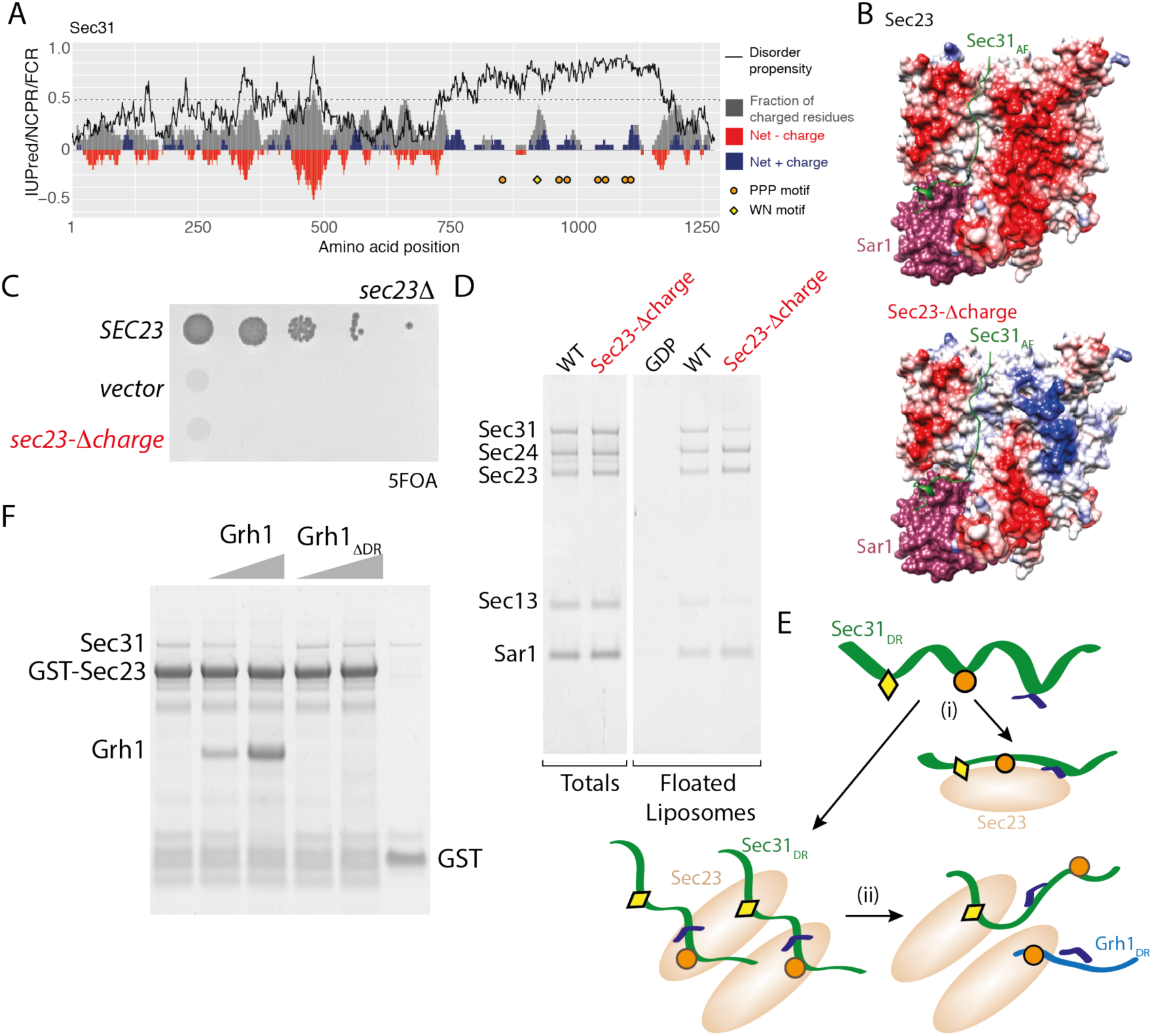
Charge interactions contribute to assembly. **A.** Charge/disorder plot for Sec31: the black curve indicates predicted disorder propensity (IUPred); a value >0.5 (dashed line) suggests intrinsic disorder. Each grey bar corresponds to fraction of charged residues in a sliding window of 20 amino acids; red/blue bars correspond to net charge per residue in a sliding window of 20 amino acids. PPP motifs are indicated by orange circles, WN motifs are indicated by yellow diamond. **B**. Surface rendering of the crystal structures of Sar1 (red) and Sec23 (coloured according to electrostatic potential) bound to the active fragment of Sec31 (green), highlighting a negatively charged patch (red region) to the right of the active fragment (left panel), which changes in electrostatic potential in the Sec23-Δcharge mutant (right panel). **C**. Serial dilutions of *sec23Δ* strains transformed with the plasmids indicated reveals lethality of charge reversal of the negative patch. **D.** Liposome flotation using the indicated proteins; recruitment of Sec13/Sec31 to liposomes coated with the Sec23-Δcharge mutant is reduced relative to wild-type. **E.** Cartoon depicting the three interface elements on the Sec31 disordered region (DR): the W_922_/N_923_ catalytic motif (yellow diamond), PPP motifs (orange circle) and charge clusters (blue cap). These elements form a multipartite binding mode such that any individual interaction can be relatively weak but still result in robust binding and oligomerization when present in combination. These elements may bind simultaneously to a single Sec23 molecule (left) or may bridge adjacent Sec23 proteins to promote an inner coat array (right).

### Disordered region-driven competition can contribute to reversible coat interactions

Using mechanistically distinct interfaces to build a global set of interactions might contribute robustness to the system while maintaining reversibility by different mechanisms: GTP hydrolysis to weaken the catalytic interface, phosphorylation of the charged regions to reverse binding, and competition from other PPP-containing proteins to disrupt the interaction interface (*12*). We sought proof of principle that a Golgi-localized PPP-containing protein might help perturb coat structure by capitalizing on the dynamic nature of an interaction interface driven by a disordered region (*17*). Grh1 is a GRASP-domain protein that together with Uso1 and Bug1 forms a COPII tethering complex that interacts with Sec23/Sec24 (*18*, *19*). The C-terminal domain of Grh1 is predicted to be disordered, with a net positively charged cluster, and a PPP motif (Fig. S4). Purified Grh1 competed for Sec31 binding to immobilized GST-Sec23, and deletion of its disordered region abrogated this effect (Fig. 3F). This observation suggests that weak PPP- and charge-driven interactions modulate coat assembly and disassembly at distinct spatial localizations (Fig. 3E(ii)).

### Coat robustness stems from combinations of interaction interfaces

Having identified multiple modes of interaction across the Sec31 disordered region, we sought to better define the importance and hierarchy of the different Sec23-binding elements. We divided the Sec31 disordered region into three segments of equivalent length and generated constructs that preserved the structural domains (the N-terminal β-propeller that forms the cage vertex, the α-solenoid rods and a C-terminal α-solenoid domain) but replaced the disordered region with different non-overlapping shortened fragments. Each segment retained at least one PPP motif and two positive-charge clusters (Fig. 4A). The first third (*sec31*_*A*_), with a single PPP motif and two charge clusters, was unable to support viability, whereas each of the two subsequent segments (*sec31*_*B*_ and *sec31*_*C*_), which both contained multiple PPP motifs and charge clusters, complemented a *sec31Δ* null strain (Fig. 4A). We note that deletion of all PPP motifs in full-length Sec31, where W/N and charge patches remain, was not lethal (Fig. 2B). In contrast, deletion of the four PPP motifs in Sec31_C_, which lacks the W/N interaction but retains charge patches, abrogated viability (Fig. 4B, *sec31*_*C*_-*ΔPPP*). This lethality suggests that PPP motifs become more important in the context of a shorter fragment that presumably has less overall affinity for Sec23. We further dissected the C-terminal portion of the disordered region into two smaller fragments, each of which contained at least two PPP motifs and one charge cluster, but neither supported viability (*sec31*_*D*_ and *sec31*_*E*_, Fig. 4A). We tested each of the shortened constructs in the liposome binding assay, observing that proteins that conferred viability also supported coat recruitment (Fig. 4C). However, in the *in vitro* budding assay, we saw distinct effects for Sec31_B_ and Sec31_C_ (Fig. 4D). Budding reactions with GTP were supported by Sec31_C_ but not Sec31_B_, consistent with the active fragment contained within Sec31_B_ stimulating GTP hydrolysis and thereby destabilizing the coat prematurely. Conversely, in the presence of GMP-PNP, Sec31_B_ supported vesicle release whereas Sec31_C_ was less effective. Again, this is consistent with the known interface occupied by the active fragment, where the Sec23/Sar1•GMP-PNP complex would stabilize association with Sec31_B_, prolonging interaction to drive vesicle formation.

**Figure 4.**
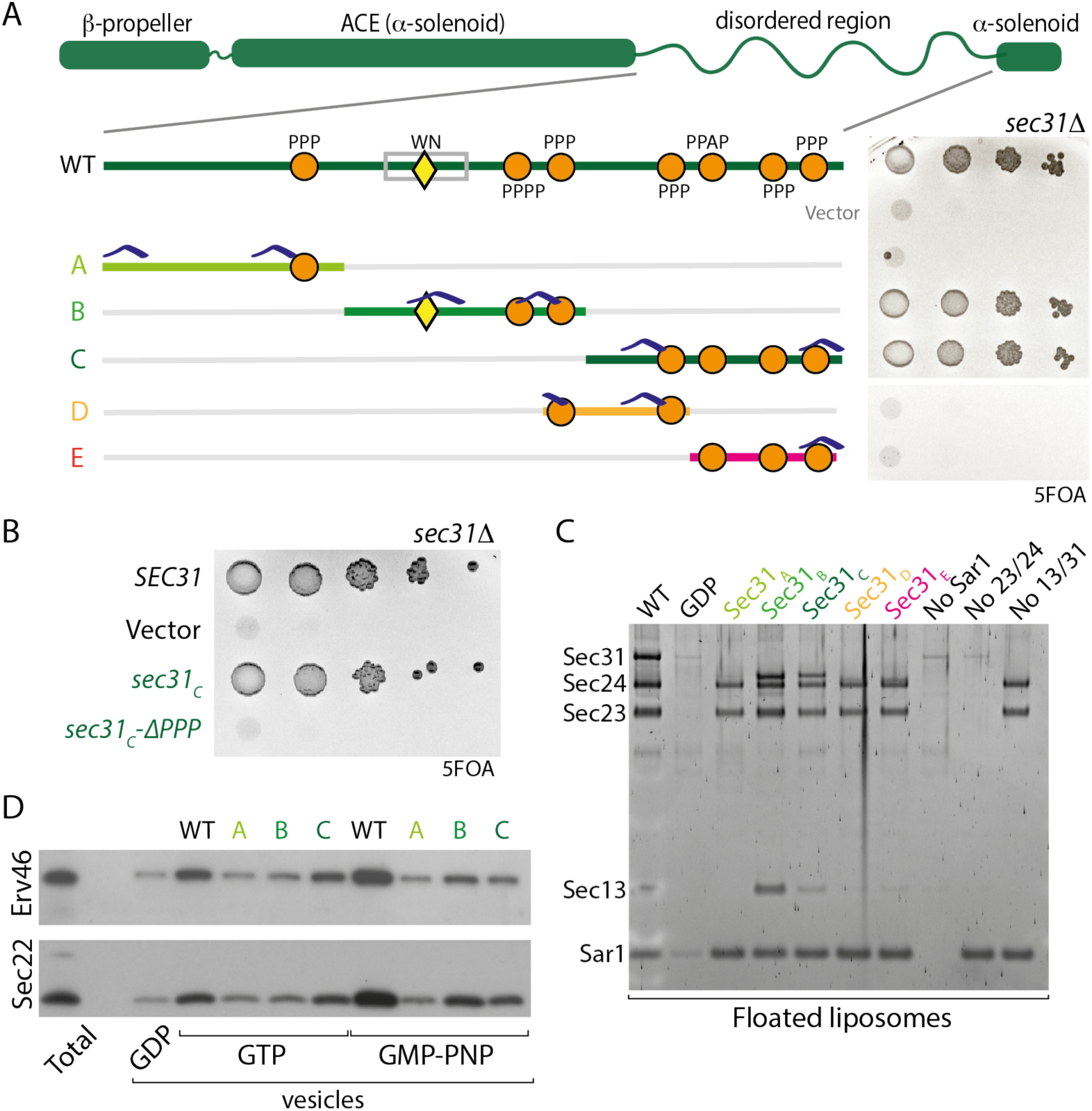
Dissection of Sec31 interaction interfaces. **A**. The disordered region of Sec31 was dissected into the indicated fragments, and the ability of these shortened regions to function in place of the full disordered region was tested by serial dilution on 5FOA; only the B and C fragments support viability. Features indicated include the W_922_/N_923_ motif (yellow triangle), PPP motifs (orange circles), and positive charge clusters (blue caps). **B**. Serial dilutions of Sec31_C_ after deletion of PPP motifs reveals the importance of these sequences in the context of a shorter disordered region. **C**. Recruitment of disordered region fragment proteins to liposomes correlates with viability: only B and C fragments conferred recruitment. **D**. *In vitro* budding from microsomal membranes with the indicated proteins also correlated with viability but showed nucleotide dependence.

The *in vitro* and *in vivo* phenotypes of these minimal constructs highlight the different assembly elements: the W/N motif that provides nucleotide-dependent affinity; linear motifs (PPP) that form weak but specific binding interfaces; a charge-driven interface; multivalency of these weak interactions; and disordered region length. The absence of the W/N interface can be compensated for by four PPP motifs and charge patches (Sec31_C_), but not by charges in the context of a single PPP (Sec31_A_). Moreover, three PPP motifs with charge patches in the context of a short fragment (Sec31_E_) cannot support assembly. Such compensation suggests that the different interfaces are partially redundant but reinforce each other to make a robust system. Orthologous Sec31 proteins across species may alter how they employ these interfaces to create functional proteins that might exhibit distinct assembly properties.

### Coat assembly can be driven by diverse sequences with conserved properties

Our charge/disorder analysis revealed conserved properties within the Sec31 disordered regions across species even though primary sequence identity was relatively low (Fig. S3B). This is in contrast to sequence conservation across the structured domains, which is relatively high (Fig. S3B), and the purified proteins show clear structural similarity. We reasoned that if coat assembly is driven by a combination of multi-modal interactions rather than protein sequence properties, a disordered region from an orthologous Sec31 might functionally replace the yeast domain. We created such chimeric proteins where the yeast disordered region was replaced with that of human Sec31A (23% similarity to the yeast Sec31 disordered region), human Sec31B (26% similarity) or Arabidopsis Sec31A (15% similarity) (Fig. 5A). Each of the substitute disordered regions, although differing in length, contained clusters of positively charged residues (Fig. S3A). Each orthologue had two PPP (or PPAP/PAPP) motifs, but both human Sec31B and Arabidopsis Sec31A lacked the catalytic W/N residues. When these constructs were introduced into yeast, only the chimera with the disordered region of HsSec31A disordered region (*sec31-Hs31A*_*DR*_) supported viability as the sole copy of Sec31 (Fig. 5A). Abrogation of the PPP motifs in this construct reduced viability (Fig. 5B), confirming that these linear motifs are indeed important in driving coat assembly. We tested the two yeast-human chimeric proteins in liposome recruitment (Fig. 5C) and budding assays (Fig. 5D), finding activity mirrored the *in vivo* phenotypes. Only Sec31-Hs31A_DR_ was able to bind Sar1/Sec23/Sec24 and to drive budding from microsomal membranes. We note that the two disordered regions that were non-functional in yeast, HsSec31B and AtSec31A, both lack catalytic W/N residues, suggesting that one of the modes of interaction is not preserved in these proteins. In both cases, these proteins are one of multiple Sec31 paralogs, suggesting that some species might employ mixed coats containing both catalytic and non-catalytic subunits. This might be a mechanism of modulating the global rate of GTPase activity, and thus coat turnover, to fine-tune coat assembly.

**Figure 5.**
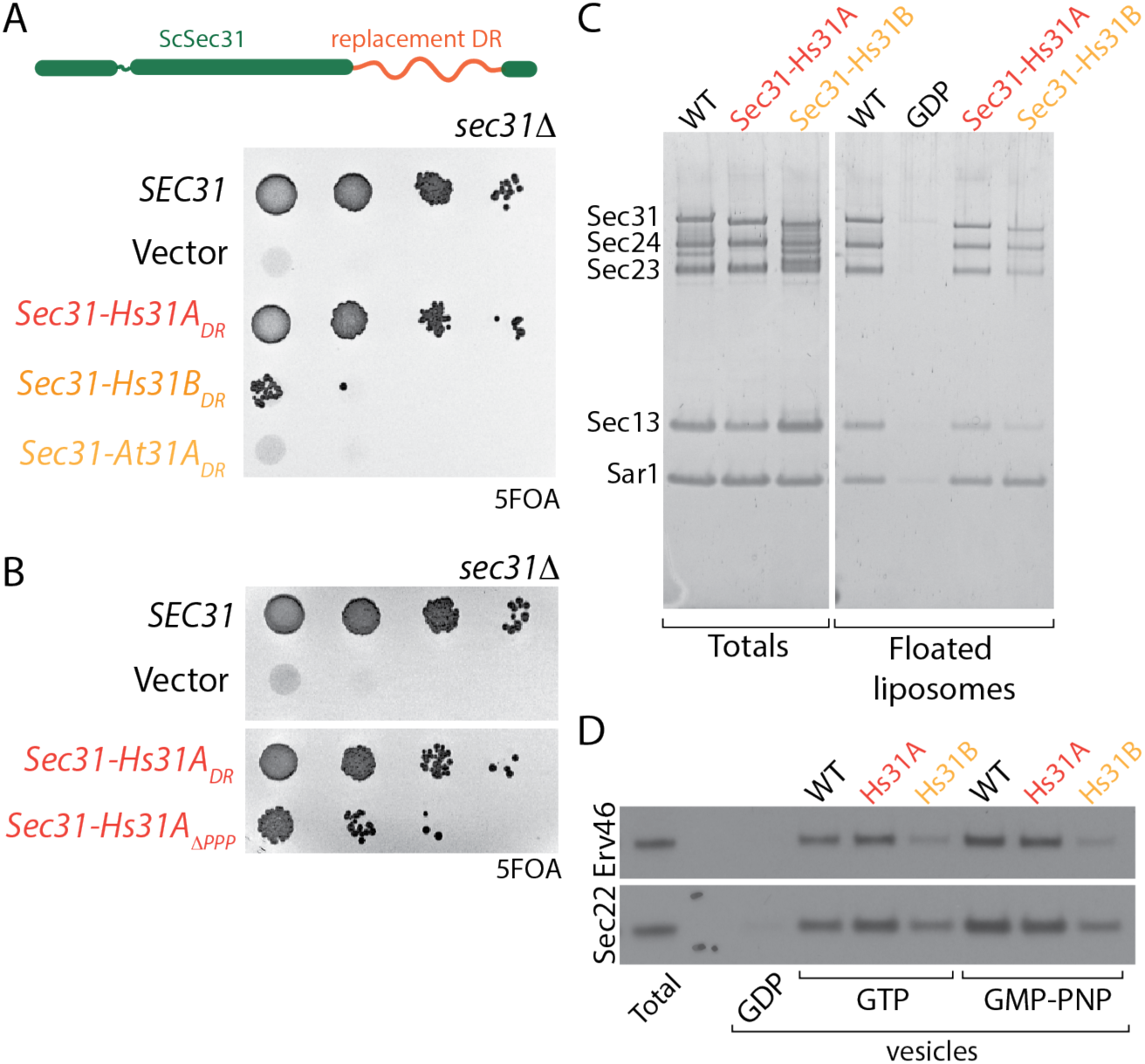
Sec31 disordered region can be functionally replaced with unrelated sequences. **A.** Serial dilutions of the indicated constructs where the yeast Sec31 disordered region was replaced with that of other species reveals that only the human Sec31A domain can support viability. **B.** Serial dilutions of the indicated strains reveals that deletion of the PPP motifs within the human Sec31A disordered region reduces viability. **C.** Liposome binding experiments with the indicated proteins correlate with viability: coat assembly is driven by the Sec31A region but not by the Sec31B region. **D.** *In vitro* microsomal budding experiments using the indicated proteins also correlate with viability: the Sec31A fusion can support vesicle formation with both GTP and GMP-PNP. **E.** Serial dilutions of chimeric Sec31 constructs containing the indicated unrelated disordered regions reveals that the Las17 domain swap supports viability.

Given the surprising finding that a human disordered region can function in yeast despite such divergence (23% similarity) in protein sequence, we sought to test even more distant domains that share similar properties. We first asked whether the disordered regions of Sec16 could similarly replace that of Sec31. Sec16 localises to ER exit sites and is known to interact with Sec23 and Sec24, likely acting as a scaffold to initiate COPII recruitment and organisation (*20*). Sec16 has two intrinsically disordered regions that lie upstream and downstream of a conserved central domain (CCD), which is a structured helical region that interacts with Sec13 (*21*) (Fig. 6A). The upstream disordered region (DR1) interacts with Sec24 (*22*) and contains a single PPP motif along with several positive charge clusters (Fig. 6A), whereas the downstream region (DR2) interacts with Sec23 (*22*), and contains multiple PPP motifs and significant areas of both net positive and negative charge (Fig. 6A). We replaced the Sec31 disordered region with Sec16_DR1_ or Sec16_DR2_ and tested each chimera for their ability to function in place of Sec31. Both Sec31-16_DR1_ and Sec31-16_DR2_ were recruited to the coat in the liposome binding assay (Fig. 6B), suggesting that each could interact with the inner coat to support initial assembly. In contrast, only Sec31-16_DR1_ could complement a *sec31Δ* strain (Fig. 6C), and support budding from microsomal membranes, albeit less efficiently than wild-type Sec31 (Fig. 6D). Since Sec31-16_DR2_ could bind to membranes but not support microsomal budding, we tested whether it could tubulate GUVs *in vitro*, and found this protein to be inactive. The discrepancy between coat binding by Sec31-16_DR2_ but failure to drive vesicle formation or support viability suggests that the mode of interaction driven by the second disordered region of Sec16 is not compatible with the specific coat conformation required for membrane bending. We note that Sec16_DR2_ has multiple features that may impact inner coat interactions, most notably significant negatively charged clusters that are not present in Sec31 disordered regions. This divergence is probably indicative of the distinct function that Sec16 serves as an accessory or regulatory factor.

**Figure 6.**
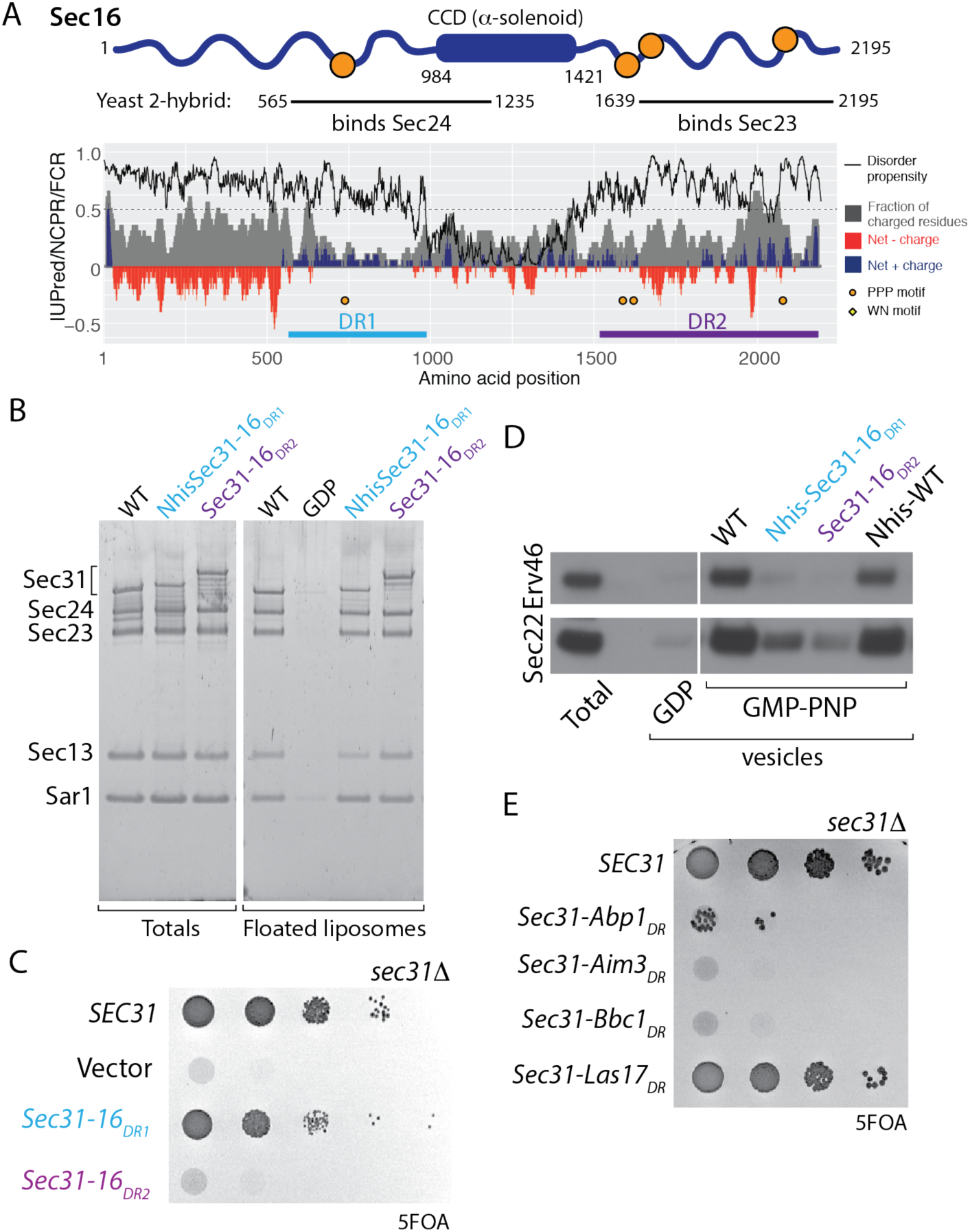
Sec16 disordered regions differentially support coat assembly. **A**. Diagram and charge-disorder plot of Sec16, highlighting the two disordered regions, DR1 (blue) and DR2 (purple) that interact with Sec24 and Sec23 respectively. Plot is annotated as described in Figure 3A. **B**. Serial dilutions of Sec31-Sec16 disordered region chimera reveals that only the first Sec16 disordered region can support viability when replacing the Sec31_DR_. **C.** Liposome binding experiments with the indicated proteins suggest that both Sec16 disordered regions confer coat recruitment. **D.** *In vitro* budding from microsomal membranes with the indicated proteins correlates with viability, albeit with reduced efficiency. **E.** Sec31 can be recruited to immobilized GST-Sec23 (lane 1), but binding is competed by excess Grh1 (lanes 2 and 3) but not by truncated Grh1 that lacks its disordered domain (lanes 4 and 5). **F.** Cartoon depicting dynamic exchange of polymeric Sec31 with a competing low-affinity interaction from the Golgi-localized tether, Grh1.

We next searched the yeast genome for additional proteins that have predicted disordered regions that also contain multiple PPP motifs. We observed that such proteins fell into three broad functional categories (Fig. S5A). The COPII category comprises Sec31 and the accessory protein, Sec16. A second category consists of proteins involved in endocytosis and actin regulation, and a third contains proteins involved in RNA processing (Fig. S5A). In a recent analysis of yeast disordered segments (*23*), several of the endocytosis/actin proteins had multiple properties (*16*) within disordered regions that were similar to Sec31 (Fig. S5B). We therefore tested whether some of these disordered regions could function in COPII coat assembly when replacing the endogenous Sec31 disordered region. Indeed, the disordered region of Las17 could functionally replace that of Sec31 to support viability (Fig. 6E). Las17 is an actin assembly factor that uses its proline rich domain to nucleate actin filaments (*24*). This mode of interaction is shared by other components of the actin/endocytic system and has parallels with the COPII coat in that multivalent weak interactions drive assembly (*25*). We note that the Las17 region we tested shares several key features with other regions that could function in the context of Sec31: multiple PPP motifs, clusters of net positive charge in the absence of significant net negative charge, and significant length. Since Las17 has no W/N motif, clearly these features alone suffice to drive coat assembly, highlighting the plasticity of the inner/outer coat interaction.

Together, our mutagenesis and disordered region replacement experiments reveal a multimodal mechanism of coat assembly, where interactions between the inner and outer coat layers comprise a multivalent fuzzy interface (*26*, *27*). Individual interactions derive from distinct elements that include linear motifs (PPP), charge (positive charge clusters), and length. Each individual interaction is dispensable, but in concert they drive function. The precise role of disordered region length remains to be explored; one possibility is that a shortened disordered region (for example the Sec31_E_ fragment which still contains multiple interaction sites) fails to assemble because of an inability to bridge adjacent Sec23 complexes. A coat assembly/disassembly mechanism driven by fuzzy interactions provides a possible mechanistic explanation for how a metastable structure like a vesicle coat can form in a controlled manner (*27*). By building up from initially weak, transient interactions, the coat can be recruited locally but is not committed to full assembly until a threshold of both inner and outer coat components is reached. Moreover, upstream accessory factors, like Sec16 and TANGO1, that use the same interactions can prime an exit site for Sec31 recruitment and organize the inner coat. Once Sec31 is engaged, its disordered region can act as molecular velcro, being strong yet readily reversible by specific mechanisms. Coat propagation could feed forward as cage vertex interactions (mediated by the β-propeller domain of Sec31) bring in additional outer coat rods to in turn organize more inner coat complexes. As the diversity of Sec31 paralogs expands, as seen in multicellular organisms, disordered regions may fuel evolution of altered interfaces that allow for fine-tuning of vesicle assembly, disassembly and geometry (*28*). For instance, the lack of the catalytic W/N motif in human Sec31B may reflect the need for a more stable coat structure, whereby the GTPase activity of the coat is not stimulated. Such stability may prolong initial events during coat assembly to favour formation of non-canonical carriers.

The multiple assembled interfaces between Sec23 and Sec31 could contribute to persistence of the polymerized coat even after GTP hydrolysis by Sar1 and its release from the membrane. Evidence for preserved coat elements on Sar1-free vesicles comes from immunogold labelling experiments that revealed persistent Sec23 and Sec13 on COPII vesicles generated with GTP (*15*). Similarly, multivalent combinatorial interactions provide an excellent mechanism for how a metastable coat polymer can be disassembled without the need for uncoating chaperones that expend energy. A combinatorial binding mode allows a stable structure to “breathe” via dynamic weak individual interactions that permit an opposing similarly weak interaction to compete and destabilize the structure. In the case of the COPII coat, long-distance tethering by extended coiled-coil proteins, like Uso1, could capture a coated vesicle, which would be brought close to the Golgi membrane, where Grh1 could locally act to initiate release of Sec13/Sec31. Such a model of locally triggered uncoating, perhaps further stimulated by Golgi-localized kinases like Hrr25 (*29*), could help ensure the directionality of vesicle traffic. Our findings of coat assembly driven by interactions involving intrinsically disordered regions are also likely to apply to other vesicle trafficking steps. Each of the major coat complexes have significant disordered regions within their subunits (*30*), and the flexible nature of the clathrin light chain is important in coat assembly (*31*). Our approach and the findings that we present here provide a conceptual framework to now discover and investigate how such elements in disordered regions of other coat complexes are used for dynamic assembly and disassembly.

## Acknowledgements

We thank Wanda Kukulski and Manu Hegde for comments on the manuscript, Jianguo Shi for help with insect cell culture, Mark McClintock for assistance with protein purification, and David Owen for advice on Sec23 mutagenesis.

## Author contributions

Conceptualization: EAM, MMB; Funding acquisition: EAM, GZ, MMB, XL; Investigation: VS, JH, GZ, XL, BS, EAM; Writing – original draft: EAM; Writing – review and editing: VS, JH, GZ, XL, BS, MMB, EAM.

## Data and materials availability

All data is available in the manuscript or the supplementary materials; plasmids and strains described can be obtained from EAM.

## Supplementary Materials

### Materials and Methods

#### Strains and plasmids

Yeast strains used in this study are listed in Table S1. Strains were constructed using standard genetic knock-out and LiAc transformation methods. Cultures were grown at 30°C in standard rich medium (YPD: 1% yeast extract, 2% peptone, and 2% glucose) or synthetic complete medium (SC: 0.67% yeast nitrogen base and 2% glucose supplemented with amino acids) as required. For testing viability, strains were grown to saturation in SC medium selecting for the mutant plasmid overnight at 30°C. 10-fold serial dilutions were made in 96 well trays before spotting onto 5FOA plates (1.675% yeast nitrogen base, 0.08% CSM, 2% glucose, 2% agar, 0.1% 5-fluoroorotic acid). Plates were scanned at day 2 or day 3 after spotting and growth at 30°C.

Plasmids used in this study are listed in Table S2. Standard cloning methods were used, including PCR amplification of yeast genes from genomic DNA, site-directed mutagenesis using the QuikChange system (Agilent) and Gibson Assembly (New England Biolabs) as per manufacturers’ instructions.

#### Protein expression and purification

Sar1 was prepared as described (*32*). Briefly, GST-Sar1 was expressed in bacterial cells by induction for 2hr with IPTG. Cells were lysed by sonication, and lysates were clarified and incubated with glutathione-agarose beads. The beads were then washed and GST-free Sar1 was generated through thrombin cleavage.

Sec23/Sec24 (Sec23 and His-Sec24) and Sec13/Sec31 (Sec13 and His-Sec31) complexes were co-expressed in Sf9 cells using the pFastbac^TM^ system. 500 ml of protein-expressing cells were collected and washed with PBS prior to freezing in liquid nitrogen. Cell pellets were lysed with a Dounce homogeniser in cold Lysis Buffer (20mM HEPES pH 8, 250mM sorbitol, 500mM KOAc, 1mM DTT, 10mM imidazole, 10% v/v glycerol). Lysates were cleared in JA 25-50 rotor (22 000 rpm, 1 hr, 4°C), and the supernatant filtered through a 0.45 μm membrane prior to loading onto a HisTrap^TM^ HP column (GE Healthcare). The following steps were done using the ÄKTA purifier system (GE Healthcare) where Elution buffer was Lysis buffer supplemented with 500 mM imidazole. The column was washed with 4% and 10% Elution buffer followed by elution using a linear gradient to 100% Elution buffer. Peak fractions were checked by SDS-PAGE followed by Coomassie staining. Verified fractions were mixed in a 3:1 ratio with QA buffer (20mM Tris pH 7.5, 1mM MgOAc, 1mM DTT, 10% v/v glycerol) and loaded onto a HiTrap^TM^ Q HP column (GE Healthcare). The protein was eluted using a linear salt gradient to a final concentration of 1M NaCl. Peak fractions were verified using SDS-PAGE and Coomassie staining and flash-frozen in liquid nitrogen. For removal of the 6xHis tag on Sec31, an overnight cleavage with His-TEV was included after the verification of Ni-IMAC fractions. The cleavage was done simultaneously with dialysis into Lysis Buffer. Uncleaved protein and His-TEV were removed by flowing through the HisTrap^TM^ HP column prior to continuing to the ion-exchange step.

6xHis-Grh1 (and the ΔDD variant) were expressed and purified as described above for Sec23/Sec24 with the following modifications: the lysates were spun in an ultracentrifuge (100, 000 x g, 1 hr, 4°C). Following the Ni-IMAC step the verified fractions were dialysed overnight in Q* buffer (20 mM Tris pH 7.5, 1 mM MgOAc, 1 mM DTT, 150 mM KOAc, 10% v/v glycerol). The sample was then concentrated using an sAmicon^®^ Centrifugal Filter Unit (MERCK) and frozen in liquid nitrogen.

#### In vitro budding from microsomal membranes

Microsomal membranes were prepared from yeast as described (*33*). Briefly, yeast cells were grown to mid-log phase in YPD (1% yeast extract, 2% peptone, and 2% glucose), harvested and resuspended in 100mM Tris pH 9.4/10mM DTT to 40 OD_600_/ml, then incubated at room temperature for 10 min. Cells were collected by centrifugation and resuspended to 40 OD_600_/ml in lyticase buffer (0.7M sorbitol, 0.75X YPD, 10mM Tris pH 7.4, 1mM DTT + lyticase 2µL/OD_600_), then incubated at 30ºC for 30 min with gentle agitation. Cells were collected by centrifugation, washed once with 2X JR buffer (0.4M sorbitol, 100mM KOAc, 4mM EDTA, 40mM HEPES pH 7.4) at 100 OD_600_/ml, then resuspended in 2X JR buffer at 400 OD_600_/ml prior to freezing at −80ºC. Spheroplasts were thawed on ice, and an equal volume of ice cold dH20 added prior to disruption with a motor-driven Potter Elvehjem homogenizer at 4ºC. The homogenate was cleared by low speed centrifugation and crude membranes collected by centrifugation of the low-speed supernatant at 27,000 x g. The membrane pellet was resuspended in ~6mL of buffer B88 (20mM HEPES pH 6.8, 250mM sorbitol, 150mM KOAc, 5mM Mg(OAc)_2_) and loaded onto a step sucrose gradient composed of 1mL 1.5M sucrose in B88 and 1mL 1.2M sucrose in B88. Gradients were subjected to ultracentrifugation at 190,000 x g for 1h at 4ºC. Microsomal membranes were collected from the 1.2M/1.5M sucrose interface, diluted 10-fold in B88 and collected by centrifugation at 27,000 x g. The microsomal pellet was resuspended in a small volume of B88 and aliquoted in 1mg total protein aliquots until use.

Budding reactions were performed as described (*6*). Briefly, 1mg of microsomal membranes per 6-8 reactions was washed 3x with 2.5 M urea in B88 and 3x with B88. Budding reactions were set up in B88 to a final volume of 250 μl at the following concentrations: 10μg/μl Sar1, 10μg/μl Sec23/Sec24, 20μg/μl Sec13/Sec31 and 0.1mM nucleotide. Where appropriate, an ATP regeneration mix was included (final concentration 1mM ATP, 50μM GDP-mannose, 40mM creatine phosphate, 200μg/ml creatine phosphokinase). Reactions were incubated for 30 min at 25°C and a 12 μl aliquot collected as the total fraction. The vesicle-containing supernatant was collected after pelleting the donor membrane (15 000 rpm, 2 min, 4°C). Vesicle fractions were then collected by centrifugation in a Beckman TLA-55 rotor (50 000 rpm, 25 min, 4°C). The supernatant was aspirated, the pelleted vesicles resuspended in SDS sample buffer and heated for 10 min at 55°C with mixing. The samples were then analysed by SDS-PAGE and immunoblotting for Sec22 (Miller lab antibody) and Erv46 (a gift from Charles Barlowe).

#### In vitro binding to small synthetic liposomes

Liposome binding experiments were performed as described (*6*). Briefly, synthetic liposomes of “major/minor” composition (50 mol% phosphatidylcholine, 21 mol% phosphatidylethanolamine, 8 mol% phosphatidylerine, 5 mol% phosphatidic acid, 9 mol% phosphatidylinositol, 2.2 mol% phosphatidylinositol-4-phosphate, 0.8% mol% phosphatidylinositol-4,5-bisphosphate, 2 mol% cytidine-diphosphate-diacylglycerol, supplemented with 2 mol% TexasRed-phosphatidylethanolamine and 20% (w/w) ergosterol) were dried to a lipid film in a rotary evaporator and rehydrated in HKM buffer (20mM HEPES pH 7.0, 160mM KOAc, 1mM MgCl_2_). The lipid suspension was extruded 17 times through a polycarbonate filter of 0.4 µm pore size. Purified COPII components and lipids were mixed to final concentrations of 0.27mM liposomes, 15μg/ml Sar1, 20μg/ml Sec23/Sec24, 30μg/ml Sec13/Sec31 and 0.1mM nucleotide in 75μl HKM Buffer. Binding reactions were incubated for 30 min at 25°C. Each sample was mixed with 50μl 2.5M Sucrose-HKM, then 100µL transferred to a ultracentrifuge tube, overlaid with 100μl 0.75M Sucrose-HKM and 20μl HKM. The gradients were spun (100 000 rpm, 25 min, 24°C with slow acceleration/deceleration) in a Beckman TLA-100 rotor. The top 30μl of the gradients were collected and normalised for lipid recovery using Typhoon FLA 7000 scanner (GE). Samples were then resolved by SDS-PAGE and visualised using SYPRO Red staining.

#### GUV budding and electron microscopy

GUVs of “major/minor” composition were made by electroformation, as described previously (*3*). Purified COPII proteins (1μM Sar1, 320nM Sec23/24 and 170nM Sec13/31) were incubated with GUVs (10% v/v), 1mM GMP-PNP (Sigma) and 2.5mM EDTA in HKM buffer (20 mM HEPES, 50 mM KOAc, 1.2 mM MgCl2, pH 6.8) at room temperature for 1-2 hours. Mutants were used in some reactions, as indicated. For negative stain electron microscopy, 4µL of the reconstitution was added to negatively glow-discharged grids (EM Sciences, CF300-Cu) and stained with 2% uranyl acetate following standard procedures. Grids were viewed with the F20 (Tecnai 200 kV) electron microscope equipped with a DE20 (Direct Electron, CA) detector. Images were collected as unaligned and summed frames with a dose of 20-30 e-/pixel/second.

#### GST-Sec23 pulldowns

For the competition assays 40μl of 50% slurry of glutathione Magbeads (GenScript) was pre-coupled with 100nM GST or GST-Sec23 for 1 hr at 4°C in binding buffer (25mM HEPES pH 7.5; 150mM NaCl; 1mM DTT; 2mM EDTA; 0.5 mM MgCl2; 2% v/v Triton X-100; protease inhibitors (Roche)). Beads were washed once with binding buffer and mixed with 500nM Sec13/Sec31 and Grh1 or Grh1-ΔDD (at 100nM or 1mM) in a total volume of 150μl and rotated for 2 hr at 4°C. Beads were washed three times with binding buffer and the samples eluted in 25μl 3x SDS Sample Buffer. Samples were heated for 5 min at 98°C, separated by SDS-PAGE and stained with SYPRO ruby.

#### Bioinformatic analyses

For analyses of sequence features of disordered domains, sequences of *Saccharomyces cerevisiae* Sec31, Sec16, Las17, Grh1, as well as human Sec31A, Sec31B, and *Arabidopsis* Sec31A were collected from Uniprot. The residue specific disorder propensity was calculated using IUPred2A using long disorder mode (*34*). Charge properties, including fraction of charged residues (FCR) and net charge per residue (NCPR), were calculated using package localCIDER (*35*), with a sliding window size of 20. PPP motifs are identified as PPP_n_ (n>2), PPAP or PAPP. The analyses were performed in Python. Data were assembled and plotted by R using custom-written scripts. For sequence similarities between disordered domains of Sec31 orthologs, yeast Sec31 (749-1174), human Sec31A (780-1112), human Sec31B (811-1078), and Arabidopsis Sec31A (722-860) were chosen for analysis. The residues were chosen based on cutoff of continuous residues predicted to have high disordered propensity (IUPred>0.5). Due to the striking difference among the lengths of these domains, three methods of alignment, pairwise local, pairwise global, multiple sequence alignment (MSA), were used to retrieve pairwise sequence similarity between the yeast Sec 31 disordered domain and each of the three homologs. Sequence alignments were performed using EMBOSS water, EMBOSS needle, Clustal Omega respectively, with default parameters (*36*).

---

**Figure S1.**
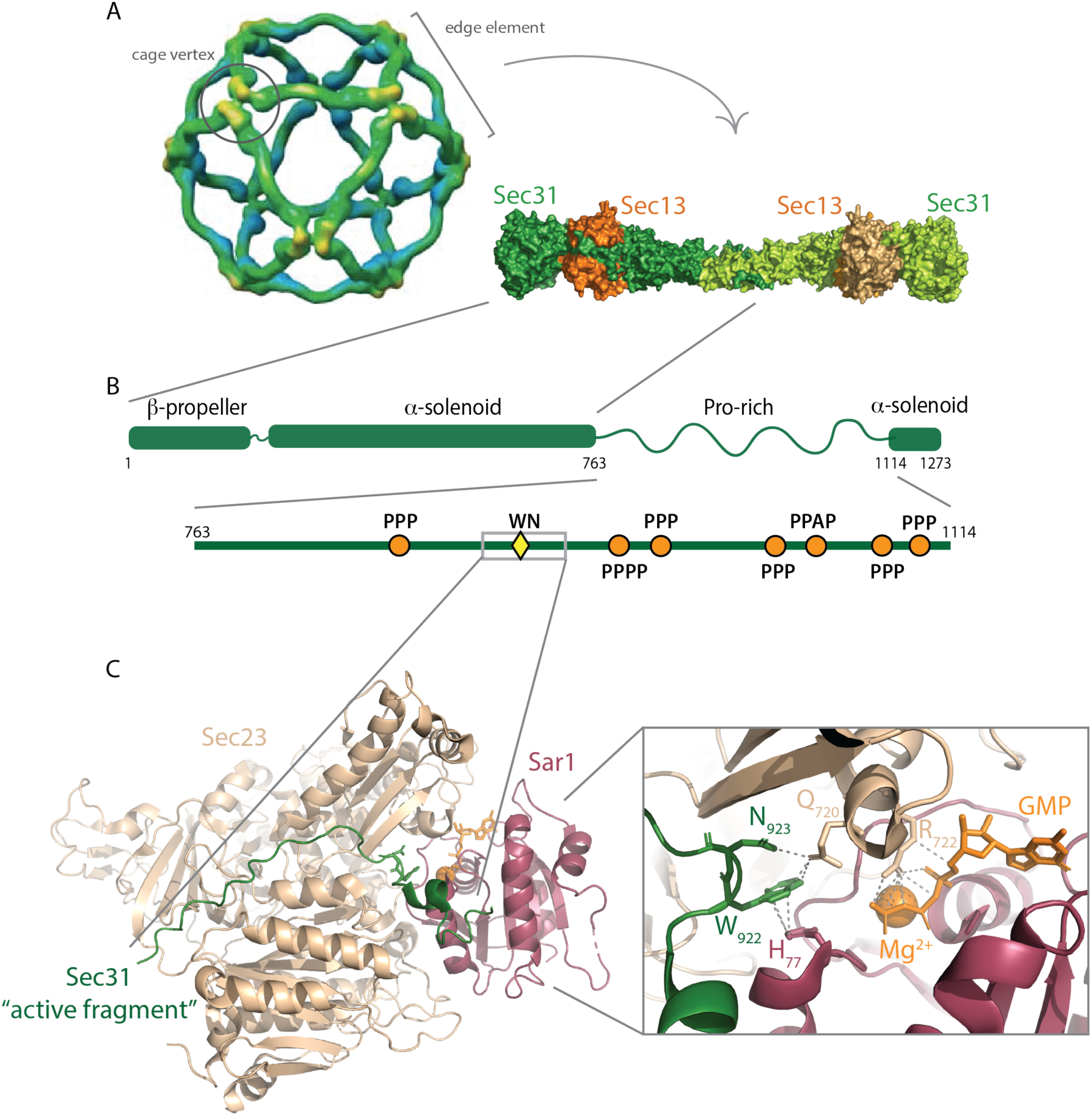
Structure and assembly of the COPII outer coat. **A.** Cryo-EM model of the cage structure formed by mammalian Sec13/Sec31 (*9*) showing the “cage vertex” where Sec31 - propeller domains tetramerize, and crystal structure (PDB 2PM6 and 2PM9 (*8*)) of the yeast Sec13/Sec31 “edge element” that forms the assembly unit (*8*). **B.** Cartoon of Sec31 showing structured elements (β-propeller and α-solenoid) and the large proline-rich disordered domain that contacts Sec23/Sar1. Within the Pro-rich domain, several motifs are noted, including triple-proline (PPP) motifs (orange circles), and the “active fragment”, which contains the key W_922_ and N_923_ residues that stimulate GTP hydrolysis on Sar1 (yellow diamond). **C.** Crystal structure (PDB 2QTV (*11*)) of Sar1/Sec23 bound to the active fragment of Sec31 (green), with GTP binding pocket enlarged (right) showing how the catalytic W/N residues help position the catalytic H_77_ and other residues within the GTP binding pocket.

**Figure S2.**
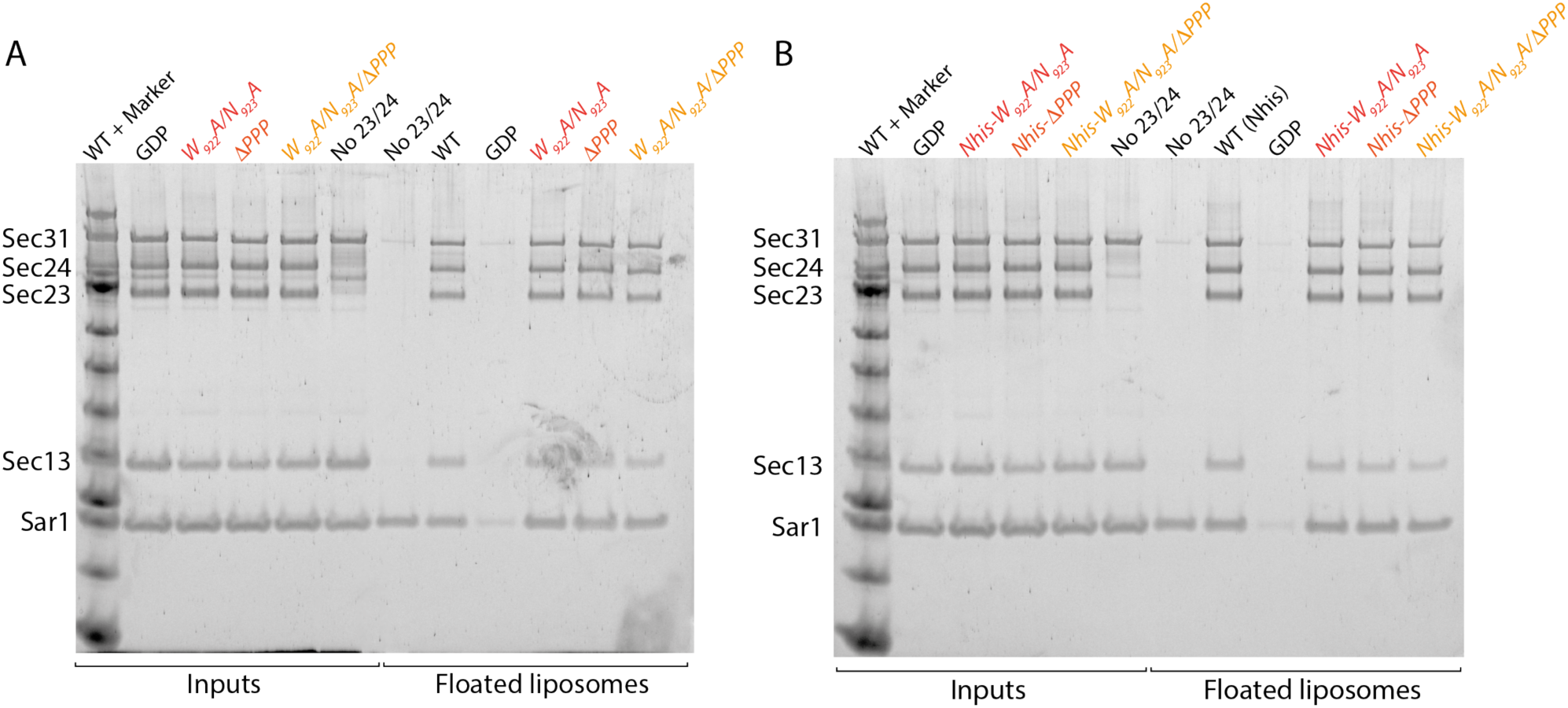
Coat assembly on synthetic liposomes is not perturbed by Sec31 mutations. Purified Sar1, Sec23/Sec24 and Sec13/Sec31 were incubated with synthetic unilamellar liposomes in the presence of GMP-PNP (inputs), and the liposomes purified by flotation through a sucrose gradient. Bound material (floated liposomes) was collected and examined by SDS-PAGE where assembled coat proteins can be distinguished. Mutation of the catalytic residues (W922A/N923A) or PPP motifs (ΔPPP) did not reduce recruitment into the coat in either the absence **(A)** or presence **(B)** of an additional N-terminal hexahistidine tag on Sec31.

**Figure S3.**
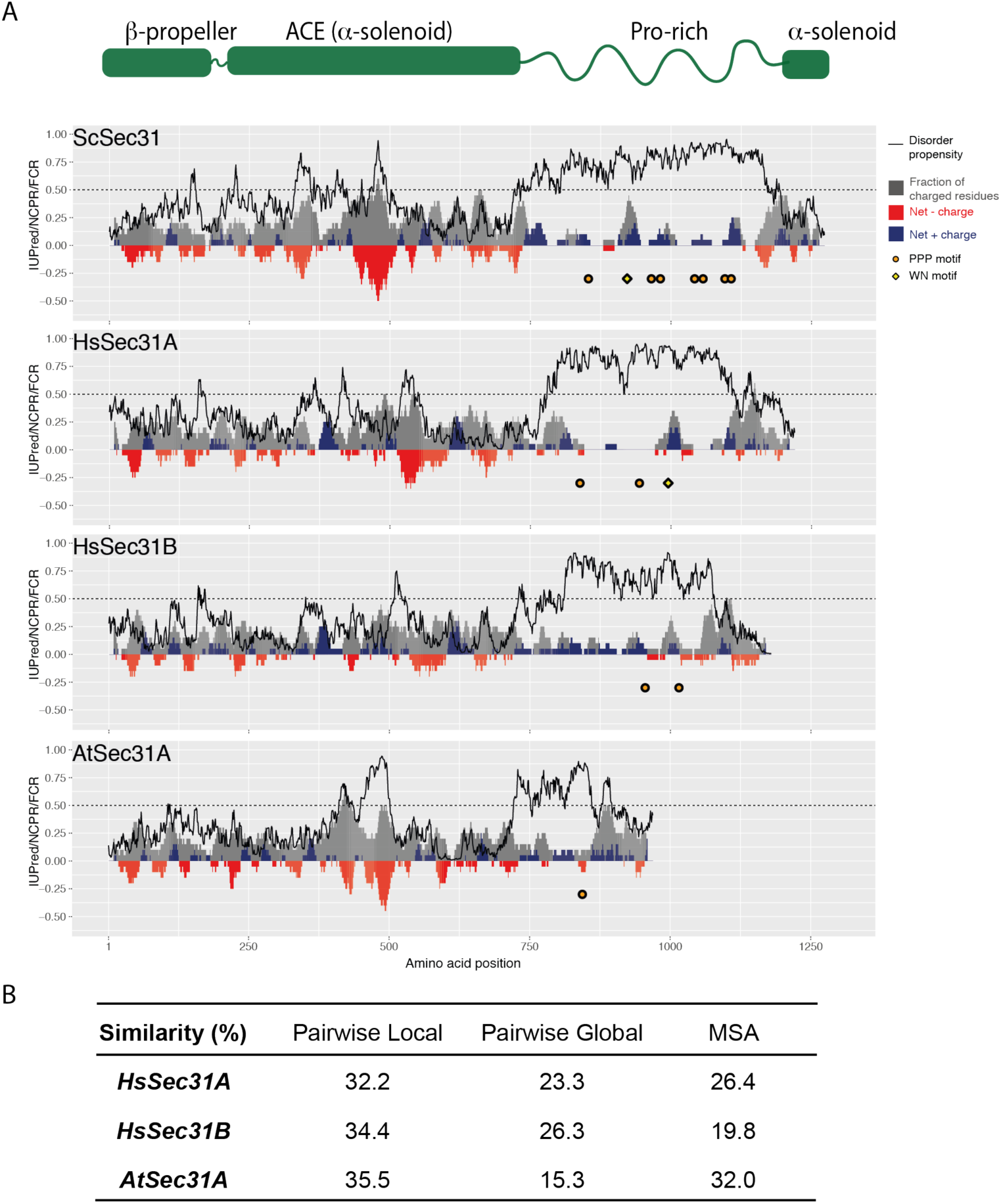
Sec31 disordered domains share several features across species. **A.** Charge-disorder plots for Sec31 orthologs from Saccharomyces cerevisiae (ScSec31), human (HsSec31A and HsSec31B) and Arabidopsis thaliana (AtSec31A). The black curve indicates predicted disorder propensity as calculated by IUPred. A value of IUPred>0.5 (dashed line) suggests a strong propensity for being intrinsically disordered. Each grey bar corresponds to fraction of charged residues (FCR) in a sliding window of 20 amino acids, centered at the residue indicated. Red/blue bars at each position correspond to net charge per residue (NCPR) in a sliding window of 20 amino acids. PPP motifs are indicated by orange circles, WN motifs are indicated by yellow diamond. Human Sec31A is the only ortholog that has a disordered domain containing three features: a WN motif, multiple PPP motifs, multiple positively charged clusters. **B.** Sequence similarity between ScSec31 disordered domains and corresponding domains in orthologs. Three methods of alignment have been used (see methods). None of the orthologs share significant similarity to ScSec31, either globally or locally.

**Figure S4.**
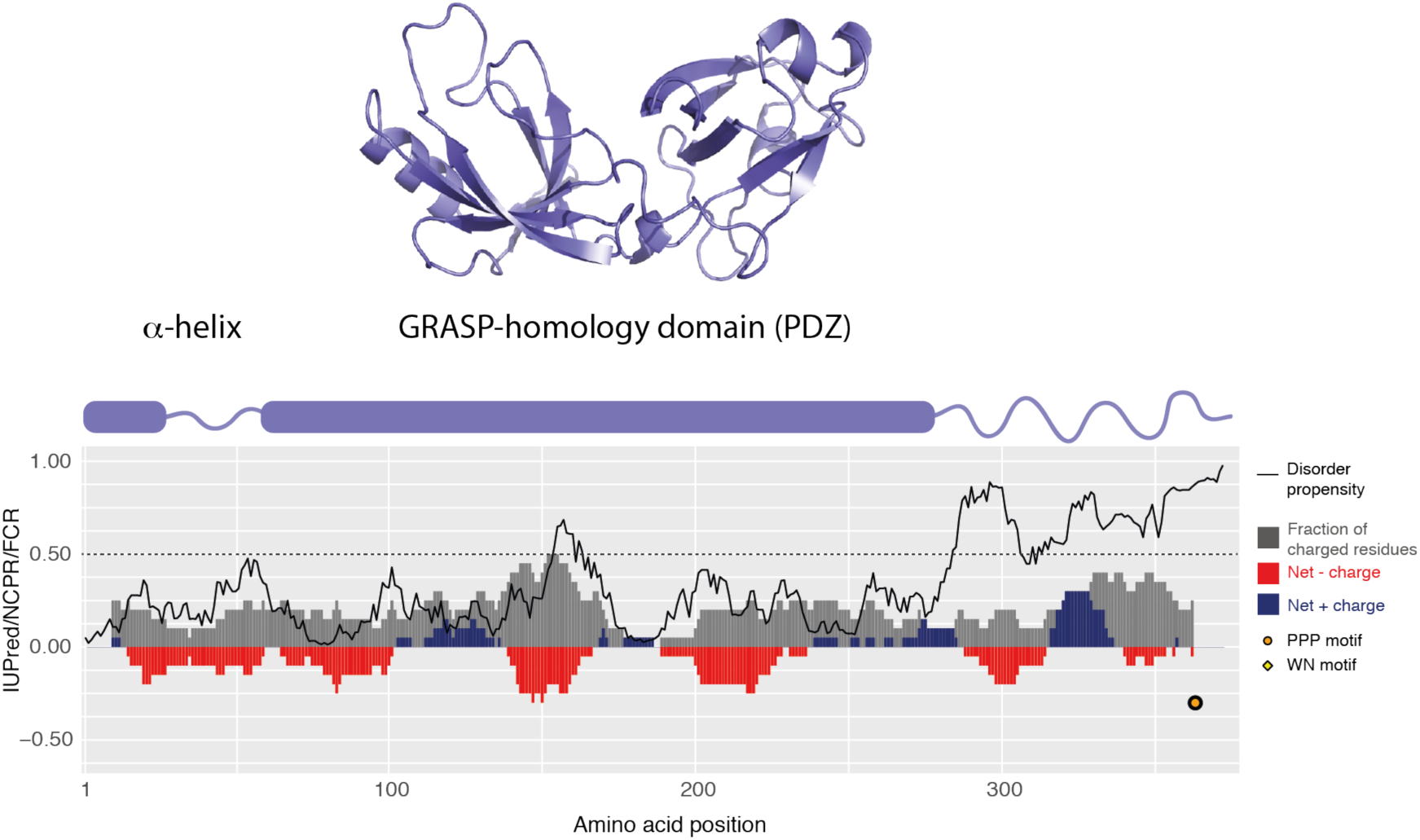
Grh1 contains a disordered domain with PPP motifs. Diagrams of Grh1 structure (blue) showing structural domains (N-terminal α-helix, and a PDZ-like GRASP homology domain), and charge-disorder along the polypeptide as described for Figure S3. The C-terminus of Grh1 is predicted to be intrinsically disordered, with a large positively charged cluster and a PPP motif.

**Figure S5.**
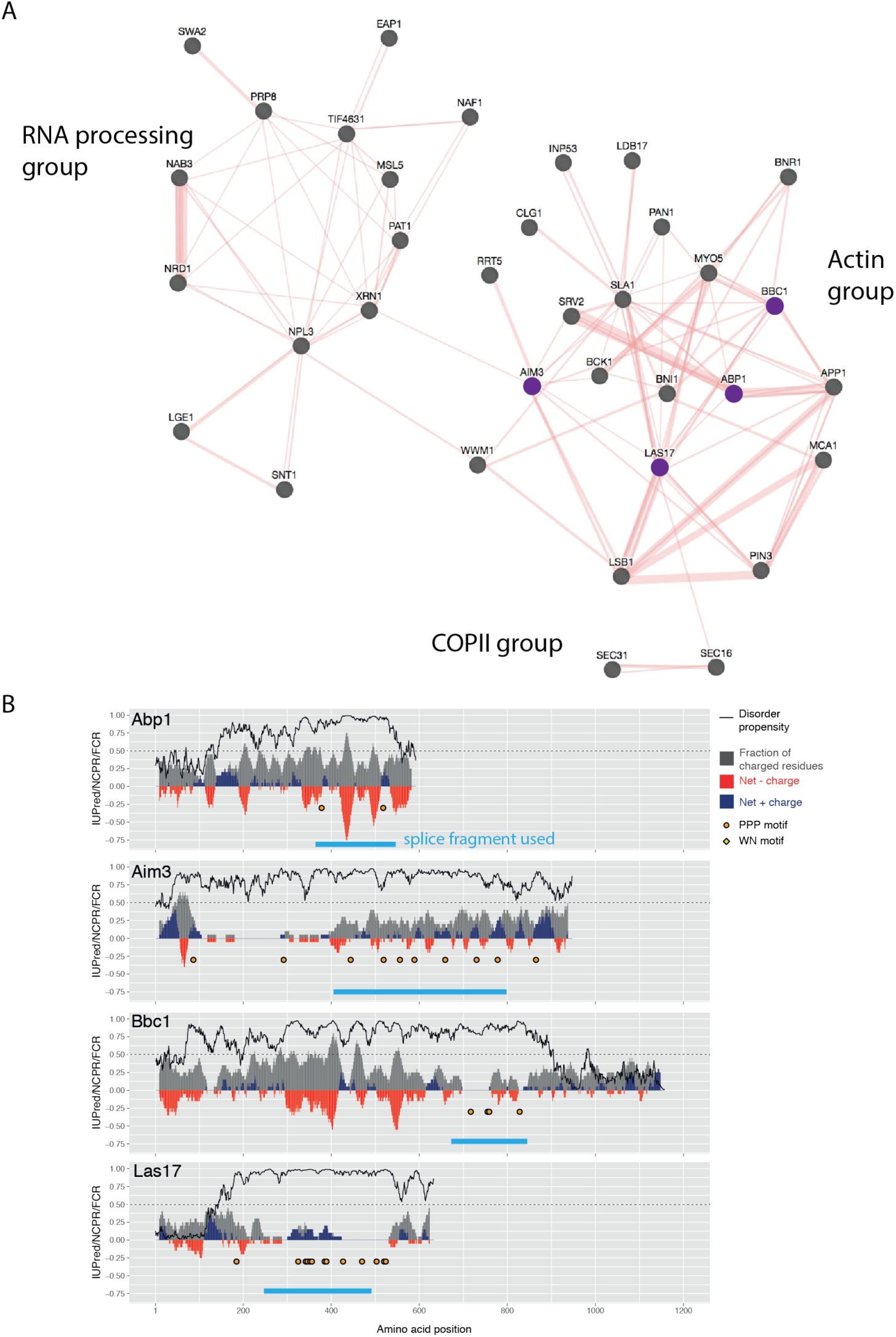
Disordered domains can be found in cytosolic proteins from multiple pathways. **A.** Protein interaction network diagram of cytosolic proteins that contain intrinsically disordered domains with multiple PPP motifs. Three functional groups are apparent: the COPII coat with Sec31 and Sec16; an endocytosis/actin module; and an RNA processing module. **B.** Charge/disorder plots for a subset of proteins identified in (A) and selected for testing as replacements for the Sec31 disordered domain. Fragments selected for splicing into Sec31 are indicated by the blue bars.

**Table S1.**
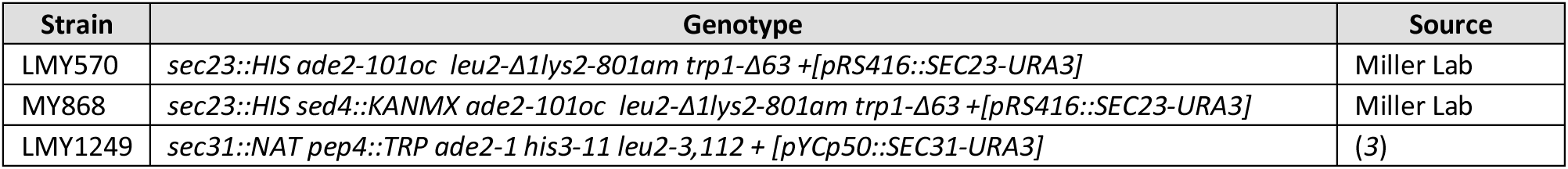
Yeast strains

| Strain | Genotype | Source |
| --- | --- | --- |
| LMY570 | *sec23::HIS ade2-101oc leu2-Δ1lys2-801am trp1-Δ63 +[pRS416::SEC23-URA3]* | Miller Lab |
| MY868 | *sec23::HIS sed4::KANMX ade2-101oc leu2-Δ1lys2-801am trp1-Δ63 +[pRS416::SEC23-URA3]* | Miller Lab |
| LMY1249 | *sec31::NAT pep4::TRP ade2-1 his3-11 leu2-3,112 + [pYCp50::SEC31-URA3]* | (3) |

**Table S2.**
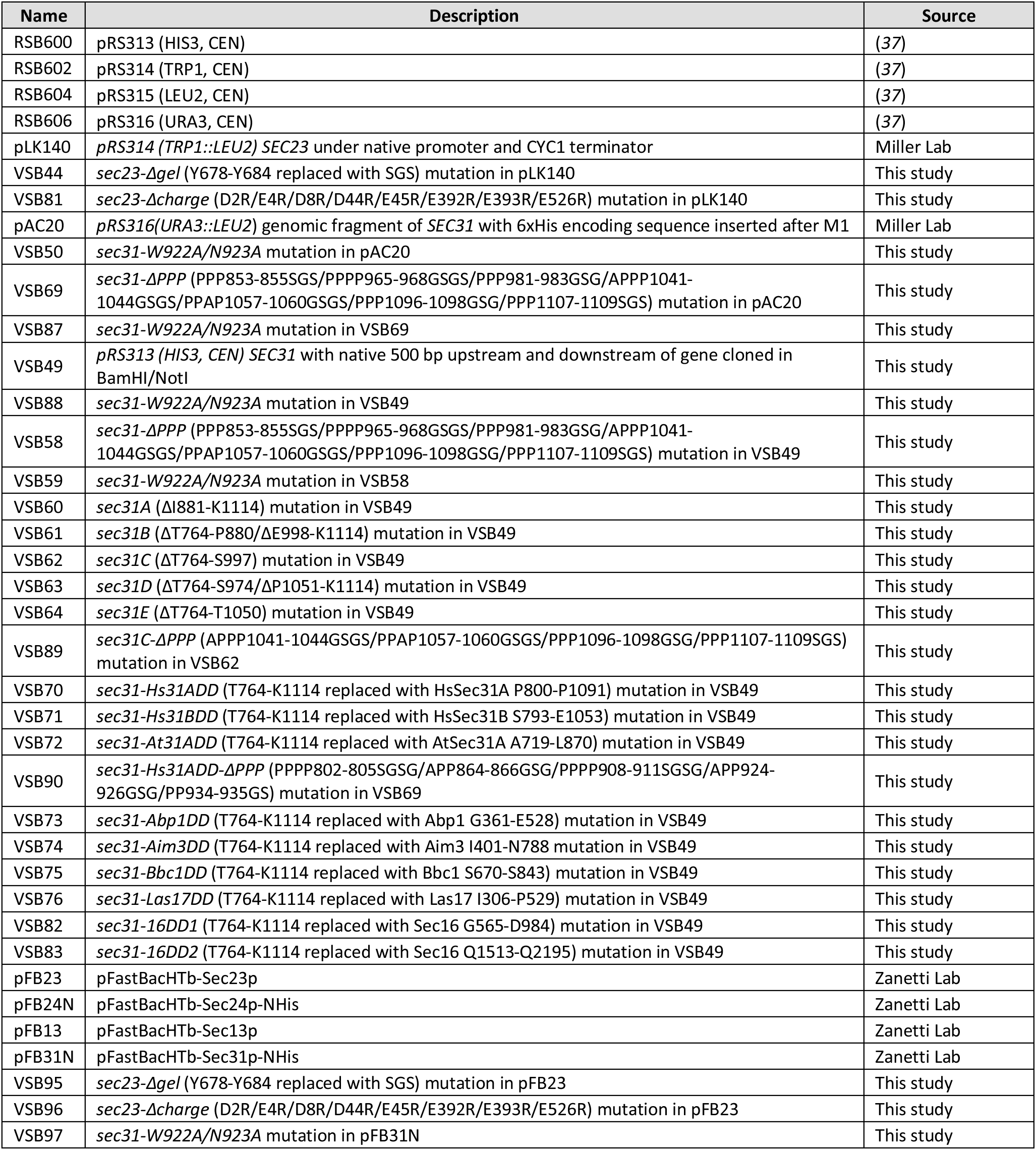
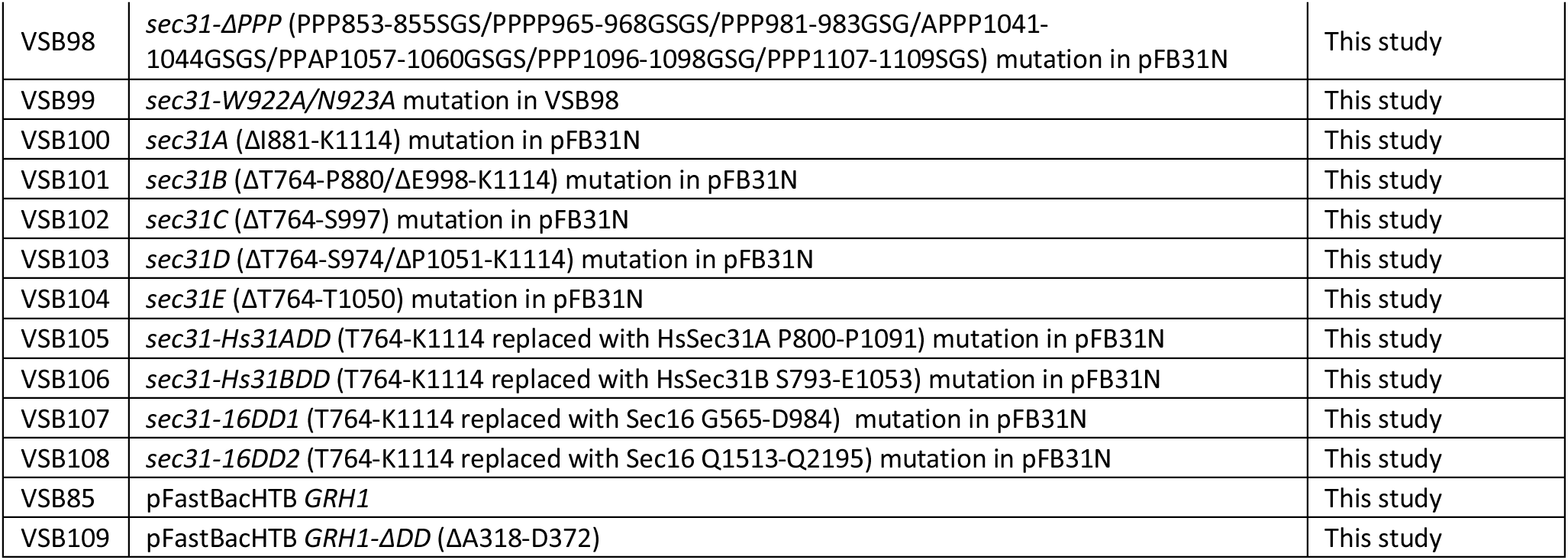
Plasmids

## References

1. J. C. Stachowiak, F. M. Brodsky, E. A. Miller, A cost-benefit analysis of the physical mechanisms of membrane curvature. Nat Cell Biol. 15, 1019–1027 (2013).

2. J. Derganc, B. Antonny, A. Copič, Membrane bending: the power of protein imbalance. Trends Biochem Sci. 38, 576–584 (2013).

3. J. Hutchings, V. Stancheva, E. A. Miller, G. Zanetti, Subtomogram averaging of COPII assemblies reveals how coat organization dictates membrane shape. Nature Communications. 9, 4154–8 (2018).

4. K. Matsuoka et al., COPII-coated vesicle formation reconstituted with purified coat proteins and chemically defined liposomes. Cell. 93, 263–275 (1998).

5. E. Mossessova, L. C. Bickford, J. Goldberg, SNARE selectivity of the COPII coat. Cell. 114, 483–495 (2003).

6. E. Miller, B. Antonny, S. Hamamoto, R. Schekman, Cargo selection into COPII vesicles is driven by the Sec24p subunit. EMBO J. 21, 6105–6113 (2002).

7. B. Antonny, D. Madden, S. Hamamoto, L. Orci, R. Schekman, Dynamics of the COPII coat with GTP and stable analogues. Nat Cell Biol. 3, 531–537 (2001).

8. S. Fath, J. D. Mancias, X. Bi, J. Goldberg, Structure and organization of coat proteins in the COPII cage. Cell. 129, 1325–1336 (2007).

9. S. M. Stagg et al., Structure of the Sec13/31 COPII coat cage. Nature. 439, 234–238 (2006).

10. G. Zanetti et al., The structure of the COPII transport-vesicle coat assembled on membranes. Elife. 2, e00951–e00951 (2013).

11. X. Bi, J. D. Mancias, J. Goldberg, Insights into COPII coat nucleation from the structure of Sec23.Sar1 complexed with the active fragment of Sec31. Dev Cell. 13, 635–645 (2007).

12. W. Ma, J. Goldberg, TANGO1/cTAGE5 receptor as a polyvalent template for assembly of large COPII coats. Proceedings of the National Academy of Sciences. 113, 10061–10066 (2016).

13. L. F. Kung et al., Sec24p and Sec16p cooperate to regulate the GTP cycle of the COPII coat. EMBO J. 31, 1014–1027 (2012).

14. R. E. Gimeno, P. Espenshade, C. A. Kaiser, SED4 encodes a yeast endoplasmic reticulum protein that binds Sec16p and participates in vesicle formation. J Cell Biol. 131, 325–338 (1995).

15. C. Barlowe et al., COPII: a membrane coat formed by Sec proteins that drive vesicle budding from the endoplasmic reticulum. Cell. 77, 895–907 (1994).

16. R. van der Lee et al., Classification of intrinsically disordered regions and proteins. Chem. Rev. 114, 6589–6631 (2014).

17. R. B. Berlow, H. J. Dyson, P. E. Wright, Hypersensitive termination of the hypoxic response by a disordered protein switch. Nature. 543, 447–451 (2017).

18. R. Behnia, F. A. Barr, J. J. Flanagan, C. Barlowe, S. Munro, The yeast orthologue of GRASP65 forms a complex with a coiled-coil protein that contributes to ER to Golgi traffic. J Cell Biol. 176, 255–261 (2007).

19. M. Schuldiner et al., Exploration of the Function and Organization of the Yeast Early Secretory Pathway through an Epistatic Miniarray Profile. Cell. 123, 507–519 (2005).

20. F. Supek, D. T. Madden, S. Hamamoto, L. Orci, R. Schekman, Sec16p potentiates the action of COPII proteins to bud transport vesicles. J Cell Biol. 158, 1029–1038 (2002).

21. J. R. R. Whittle, T. U. Schwartz, Structure of the Sec13-Sec16 edge element, a template for assembly of the COPII vesicle coat. J Cell Biol. 190, 347–361 (2010).

22. R. E. Gimeno, P. Espenshade, C. A. Kaiser, COPII coat subunit interactions: Sec24p and Sec23p bind to adjacent regions of Sec16p. Mol Biol Cell. 7, 1815–1823 (1996).

23. T. Zarin et al., Proteome-wide signatures of function in highly diverged intrinsically disordered regions. Elife. 8, 1727 (2019).

24. A. N. Urbanek, A. P. Smith, E. G. Allwood, W. I. Booth, K. R. Ayscough, A Novel Actin-Binding Motif in Las17/WASP Nucleates Actin Filaments Independently of Arp2/3. Curr Biol. 23, 196–203 (2013).

25. Y. Sun et al., Switch-like Arp2/3 activation upon WASP and WIP recruitment to an apparent threshold level by multivalent linker proteins in vivo. Elife. 6, 60 (2017).

26. M. Fuxreiter, Fuzziness in Protein Interactions—A Historical Perspective. Journal of Molecular Biology. 430, 2278–2287 (2018).

27. H. Wu, M. Fuxreiter, The Structure and Dynamics of Higher-Order Assemblies: Amyloids, Signalosomes, and Granules. Cell. 165, 1055–1066 (2016).

28. M. M. Babu, R. W. Kriwacki, R. V. Pappu, Structural biology. Versatility from protein disorder. Science. 337, 1460–1461 (2012).

29. C. Lord et al., Sequential interactions with Sec23 control the direction of vesicle traffic. Nature. 473, 181–186 (2011).

30. N. Pietrosemoli, R. Pancsa, P. Tompa, Structural Disorder Provides Increased Adaptability for Vesicle Trafficking Pathways. PLoS Comput Biol. 9, e1003144–19 (2013).

31. J. D. Wilbur et al., Conformation switching of clathrin light chain regulates clathrin lattice assembly. Dev Cell. 18, 841–848 (2010).

32. Y. Shimoni, R. Schekman, Vesicle budding from endoplasmic reticulum. Meth Enzymol. 351, 258–278 (2002).

33. L. J. Wuestehube, R. S. M. I. enzymology, 1992, [13] Reconstitution of transport from endoplasmic reticulum to golgi complex using endoplasmic reticulum-enriched membrane fraction from yeast. Elsevier

34. B. Mészáros, G. Erdős, Z. Dosztányi, IUPred2A: context-dependent prediction of protein disorder as a function of redox state and protein binding. Nucleic Acids Research. 46, W329–W337 (2018).

35. A. S. Holehouse, R. K. Das, J. N. Ahad, M. O. G. Richardson, R. V. Pappu, CIDER: Resources to Analyze Sequence-Ensemble Relationships of Intrinsically Disordered Proteins. Biophys J. 112, 16–21 (2017).

36. F. Madeira et al., The EMBL-EBI search and sequence analysis tools APIs in 2019. Nucleic Acids Research. 47, W636–W641 (2019).

37. R. Sikorski, P. Hieter, A system of shuttle vectors and yeast host strains designed for efficient manipulation of DNA in Saccharomyces cerevisiae. Genetics. 122, 19–27 (1989).

